# Holo-Hilbert Spectral-based Noise Removal Method for EEG High-Frequency Bands

**DOI:** 10.1101/2021.04.28.440961

**Authors:** Narges Moradi, Pierre LeVan, Burak Akin, Bradley G. Goodyear, Roberto C. Sotero

**Author notes:** Corresponding authors: N. Moradi and R. C. Sotero 3330 Hospital Drive NW, HSC Building, Calgary, AB, T2N 4N1, Canada. Declarations of interest: none.

## Abstract

Simultaneous EEG-fMRI is a growing and promising field, as it has great potential to further our understanding of the spatiotemporal dynamics of brain function in health and disease. In particular, there is much interest in understanding the fMRI correlates of brain activity in the gamma band (*>* 30 Hz), as these frequencies are thought to be associated with cognitive processes involving perception, attention, and memory, as well as with disorders such as schizophrenia and autism. However, progress in this area has been limited due to issues such as MR-induced artifacts in EEG recordings, which seem to be more problematic for gamma frequencies. This paper presents a noise removal method for the gamma band of EEG that is based on the Holo-Hilbert spectral analysis (HHSA), but with a new implementation strategy. HHSA uses a nested empirical mode decomposition (EMD) to identify amplitude and frequency modulations (AM and FM, respectively) by averaging over frequencies with high and significant powers. Our method examines gamma band by applying two layers of EMD to the FM and AM components, removing components with very low power based on the power-instantaneous frequency spectrum, and subsequently reconstructs the denoised gamma-band signal from the remaining components. Simulations demonstrate that our proposed method efficiently reduces artifacts while preserving the original gamma signal which is especially critical for simultaneous EEG/fMRI studies.

## I. Introduction

Electroencephalography (EEG) is a widely used technique that records electrical activity of the brain non-invasively with high temporal resolution (on the order of milliseconds) [1, 2]. EEG data has a low spatial resolution (on the order of centimeters) [1, 2]. On the other hand, functional magnetic resonance imaging (fMRI) based on the hemodynamic bloodoxygenation-level-dependent (BOLD) response localizes brain activity with a high spatial resolution (on the order of millimeters), but low temporal resolution (typically 2–3 seconds) [1, 3].

Fortunately, BOLD fMRI and EEG can be collected simultaneously to permit both high spatial and temporal resolution, which improves our ability to understand the coupling that exists between the brain’s electrical activity and hemodynamic responses, i.e., neurovascular coupling [4–9]. EEG data contains oscillations with different frequencies. BOLD signal is associated with neuronal synchronization across a number of EEG frequency bands. The powers of the alpha (8–12 Hz) and beta (12–30 Hz) bands correlate negatively with BOLD signal’s magnitude, while the gamma band (*>* 30 Hz) power correlates positively with BOLD signal’s magnitude [10–12]. It is believed that gamma activity is most directly associated with neurovascular coupling [12–20], and some studies have shown that gamma-band frequencies are the carrier frequencies associated with the temporal synchronization of intrinsic BOLD fluctuations within resting-state functional networks [12, 17, 21, 22]. Thus, the gamma band is of utmost importance in the analysis of EEG-fMRI data which provides crucial information with a high spatial and temporal resolution for brain-network studies to clinical investigations [23–26]. Low-frequency bands, like alpha and delta (0.1–4 Hz), have been vastly investigated, while EEG data in high-frequency ranges, like gamma band, is not studied as much as the low-frequency bands [27–31]. In this paper, we propose a denoising method that applies to all EEG bands. However, we focus on our method’s performance on the EEG gamma band due to the gamma-band’s importance in brain-function studies and the lack of efficient denoising methods for high-frequency EEG bands like the gamma band.

EEG simultaneously recorded during fMRI contains artifacts that cannot be simply filtered [10, 32–34]. These artifacts are primarily due to electric noise induced by the on and off switching of the magnetic field gradients [32–34]. The shape and amplitude of artifacts vary depending on the EEG electrode’s location, head motion, and the positioning of the leads and connectors [32, 33, 35]. In addition, Electromyographic (EMG) recordings of muscle contractions, recorded as part of the EEG acquisition, contain artifacts over a wide frequency range and are maximum at frequencies exceeding 30 Hz, which is within the gamma band [10, 31, 32]. Several noise removal methods have been proposed, including those that use Kalman filtering [36], the wavelet transform [37], blind source separation [38], principal component analysis (PCA) [38], and independent component analysis (ICA) [10, 39]. Most methods have concentrated on improving EEG signals at low frequencies (*<* 30 Hz) [27–29, 31], and these denoising strategies are shown to be inadequate for the gamma band [10]. Therefore, an effective noise-removal method for the gamma band is sorely needed. Of note, EEG signals are nonlinear processes with both amplitude and frequency modulations formed by linear additive or nonlinear multiplicative processes. Thus spectral-analysis techniques like Fourier [9, 37], wavelet [37] and Hilbert–Huang transform (HHT) [40] that are limited to the analysis of stationary and linear signals, and require a priori or a posteriori knowledge of the underlying physiology are not qualified and efficient to extract information from the nonlinear processes [41, 42]. Requiring a priori or a posteriori basis places restrictions on automated techniques. The previous Holo-Hilbert spectrum (HHS) method is effective in examining frequency modulated signals; however, it does not provide the frequency-domain information on the amplitude function and ignores the role of amplitude modulation [41, 43, 44].

The recently introduced Holo-Hilbert spectral-based analysis (HHSA) technique [41] examines both the amplitude modulation (AM) and frequency modulation (FM) variations of a signal simultaneously. HHSA constructs a multidimensional power spectrum that accommodates all possible temporal properties and interactions within a signal: additive and multiplicative, intra-mode (interaction within each mode) and inter-mode (interaction between modes), stationary and nonstationary, linear and nonlinear interactions. HHSA identifies the AM and FM within a signal based on the constructed power spectrum [7, 41]. According to this method, the signal of interest is first decomposed into intrinsic mode functions (IMFs) using the EMD method, known as the first layer. Then the amplitude envelope of each of the IMFs is decomposed into a set of amplitude-modulated IMFs, denoted by IMFs^AM^, constituting the second layer [7, 41]. The intra-mode frequency variations are examined within each frequency-modulated IMFs, denoted by IMFs^FM^ [45]. On the other hand, the AM frequency represents the slow-changing inter-mode frequency variations; it can be used to analyze interactions between IMFs^FM^ [7]. Although the HHSA method can go through multiple iterations until it bears no more cyclic characteristics, only two iterations are performed in practice [7, 41, 43, 46–48].

The HHSA method has been used in a number of EEG applications, including the examination of oscillatory brain activity and interactions amongst EEG frequency bands (alpha and beta bands) [7], the study of human visual-system processing [46], and the derivation of time-frequency spectrum matrices to characterize sleep stages [48]. However, previous studies applied two layers of EMD to the AM component of a signal and not to the FM component. We show that applying the second layer of the EMD method to the FM components of the signal along with AM components decomposition augments the capability of analyzing intra-mode nonlinearity. Moreover, there is no need for applying more than two layers of EMD decomposition and multiple iterations. Consequently, the computational burden would be much lesser than the HHSA method.

In the proposed method, the EEG signal is first decomposed into IMFs. These IMFs are combined to roughly correspond to the known EEG bands (e.g., gamma, beta) [5]. After constructing the gamma band, the second layer of decomposition is applied to the AM and FM components of the gamma band to compute its IMFs^AM^ and IMFs^FM^, respectively. Analogous to AM frequency identification of the HHSA method, the carrier frequency of a signal is where power is significant and high in the power-instantaneous frequency spectrum [7, 41]. Therefore, IMFs^AM^ and IMFs^FM^ with frequencies of low power are considered as a noise. Accordingly, the power spectrum of the IMFs^AM^ and IMFs^FM^ are used to identify and remove noise-related IMFs from the AM and FM components of the signal, respectively.

Another difference between the proposed and the HHSA methods is how the signal’s AM and FM components are computed. In the HHSA method, a signal is constructed based on the FM and AM frequencies with the significant- and highenergy-density; however, the signal in the proposed method is constructed by integrating the remaining IMFs^AM^ and IMFs^FM^ after removing the low power IMFs^AM^ and IMFs^FM^.

Our method enhances the determination of nonlinear interactions within the signal by providing frequency-domain information on the signal’s AM and FM components. Furthermore, compared to the HHSA method, denoising a signal by removing its noisy components is more reliable than making the denoised signal based on the specific AM and FM frequencies. This is because neural signals interaction is highly complicated. Consequently, the HHSA method will likely ignore some of the components of an EEG signal that are neural-based and informative about neural interactions. With these theorized improvements, we hypothesize that our proposed method generates more efficient and accurate gamma-band signals than the HHSA method.

## II. Methods

### A. EEG data acquisition

Resting-state EEG data were recorded using a 256-channel MR-compatible EEG system (Geodesic EEG System 400, Electrical Geodesic, Inc., Eugene, OR) from a subject with 1000 Hz sample rate for approximately 5 min [49]. Additional ECG leads were positioned on the subject’s chest. EEG was recorded simultaneously with fMRI, and channels were referenced to the vertex electrode, Cz. The cap diameter was adjusted to the subject before recording. Channels were connected to the scalp through a hypoallergenic sponge soaked in a saline solution of Potassium Chloride (KCl) and detergent. The signals were downsampled to 200 Hz, and the gradient and ballistocardiographic artifacts were removed using the methods described in [35, 50]. The subject gave informed consent in accordance with the study protocol approved by the Ethics Commission of the University of Freiburg.

### B. Data simulation

The background activity of the brain was modelled using 1/f-activity (or pink noise), and sensor noise was modelled at the channel level as additive white gaussian noise. To simulate the EEG’s gamma band, white gaussian and pink noise were both filtered between 30 and 80 Hz. The filtered noises were then added to a 30 Hz sinusoidal signal and a combination of three 30 Hz, 60 Hz and 75 Hz sinusoidal signals to yield different levels of SNR [51, 52]. The duration of the simulated signals was 50 s and the sampling rate was 200 Hz.

### C. Denoising EEG signals using the proposed method

To denoise the gamma band using the proposed approach, the following steps were followed:

1) The EEG signal was first decomposed into its IMFs using the EMD method [16]. The obtained IMFs roughly correspond to the known EEG bands [5], but if there was a case where several IMFs lie within the same band, they were added to obtain only one component per band. The main goal of this paper is denoising the gamma band; thus, the proposed method is applied only to the gamma band, which is denoted by *γ*. The gamma band signal can be written in terms of its AM and FM components as:

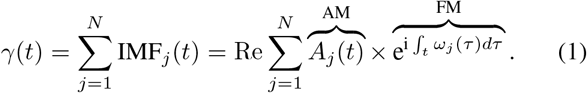

This equation corresponds to the first layer of the HHSA method, where *N* is the number of points in the time series, *A*_*j*_(*t*) are the instantaneous amplitudes, and *ω*_*j*_(*t*) are the instantaneous frequencies.

2) To extract *A*_*j*_(*t*), the absolute value of *γ*(*t*) is used to construct the upper envelope function by a natural spline function through all the maxima.

3) The *A*_*j*_(*t*) envelopes are decomposed into amplitude-modulated IMFs using the EMD method, denoted as IMF^AM^:

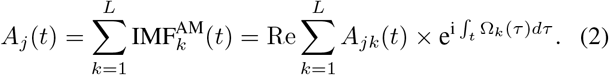

4) To have the temporal variations of the FM’s and IMF^AM^’s frequencies, the instantaneous frequencies *ω*_*j*_ and Ω_*k*_ of the IMF^AM^ are computed using the Hilbert transform [40] in the form of a time–frequency representation.

5) A three-dimensional HHSA spectrum (consisting of *ω*, Ω, and *P* (*ω*, Ω)) is constructed by incorporating the instantaneous power, *P* (*ω*, Ω), with the instantaneous AM frequencies of IMF^AM^ (Ω) and the instantaneous FM frequencies (*ω*) across all time points for the gamma band signal. The time variable is integrated out to give a frequency-only spectrum.

6) According to the HHSA method [41], the carrier-wave and envelope-modulation frequencies (*ω*, Ω) are identified based on the power density in the HHSA spectrum. The prominent modulating frequencies of the signal are where the power is high and significant and are determined by averaging over the frequencies with high and significant powers [41]. With computed *ω* and Ω, the denoised signal is constructed by Eq. (1).

The steps presented above denoise a signal according to the HHSA method. In the proposed method, the first five steps are implemented similar to the HHSA method, but the 6th step is skipped. Thus, after step 5, artifact removal analysis is performed based on the proposed method as follows:

5) *→*7) A 2D power spectrum is computed based on the instantaneous power *P* (Ω) and frequencies, Ω, across all IMFs^AM^.

8) The AM component of the signal is refined based on its 2D power spectrum, which is consistent with the HHSA spectrum but is more informative about frequencies’ power density of the IMFs. 2D power spectrum makes it easier to decide which IMFs or part of IMFs need to be removed. IMFs^AM^, or part of an IMF^AM^, of specific frequency bands are removed according to the 2D power spectrum where power is not around its peak. In other words, frequencies that are not in the bandwidth of the AM component are removed. Bandwidth is the frequency range occupied by a modulated carrier signal.

9) The FM component of the gamma-band signal is computed using the Hilbert transform.

10) The FM component, which is represented in Eq. (1) as 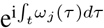, is decomposed into a set of frequency-modulated IMFs with instantaneous carrier-wave frequencies *ν*_*k*_(*τ*) using the EMD-based method, denoted as IMF^FM^.

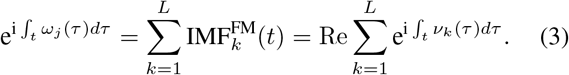

11) The instantaneous frequencies *ν*_*k*_ of IMF^FM^ are then calculated using the Hilbert Transform.

12) A 2D power spectrum of the FM component of the signal is computed based on the instantaneous power, *P*(*ν*), and frequencies, *ν*, of all IMFs^FM^.

13) To denoise the signal’s FM component, IMFs^FM^ or part of an IMF^FM^ (i. e., just some of the frequencies in a IMF) that do not have high-power are removed.

14) The remained IMFs^AM^ and IMFs^FM^ are summed up to construct the denoised AM and FM components of the denoised signal, respectively.

After processing both the AM and FM components of the signal, as shown in Eqs. (2) and (3), the integration of both the denoised AM and FM parts (*A*_*j*_(*t*)_*R*_ and 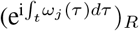 respectively), constructs the denoised gamma signal (*γ*_*R*_) as

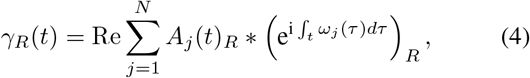

where R means noise-reduced.

The HHSA method and our proposed method were applied to the simulated datasets. A flowchart representing the overview of applying the proposed approach to denoise the gamma band signal is shown in Fig. 1.

**Fig. 1:**
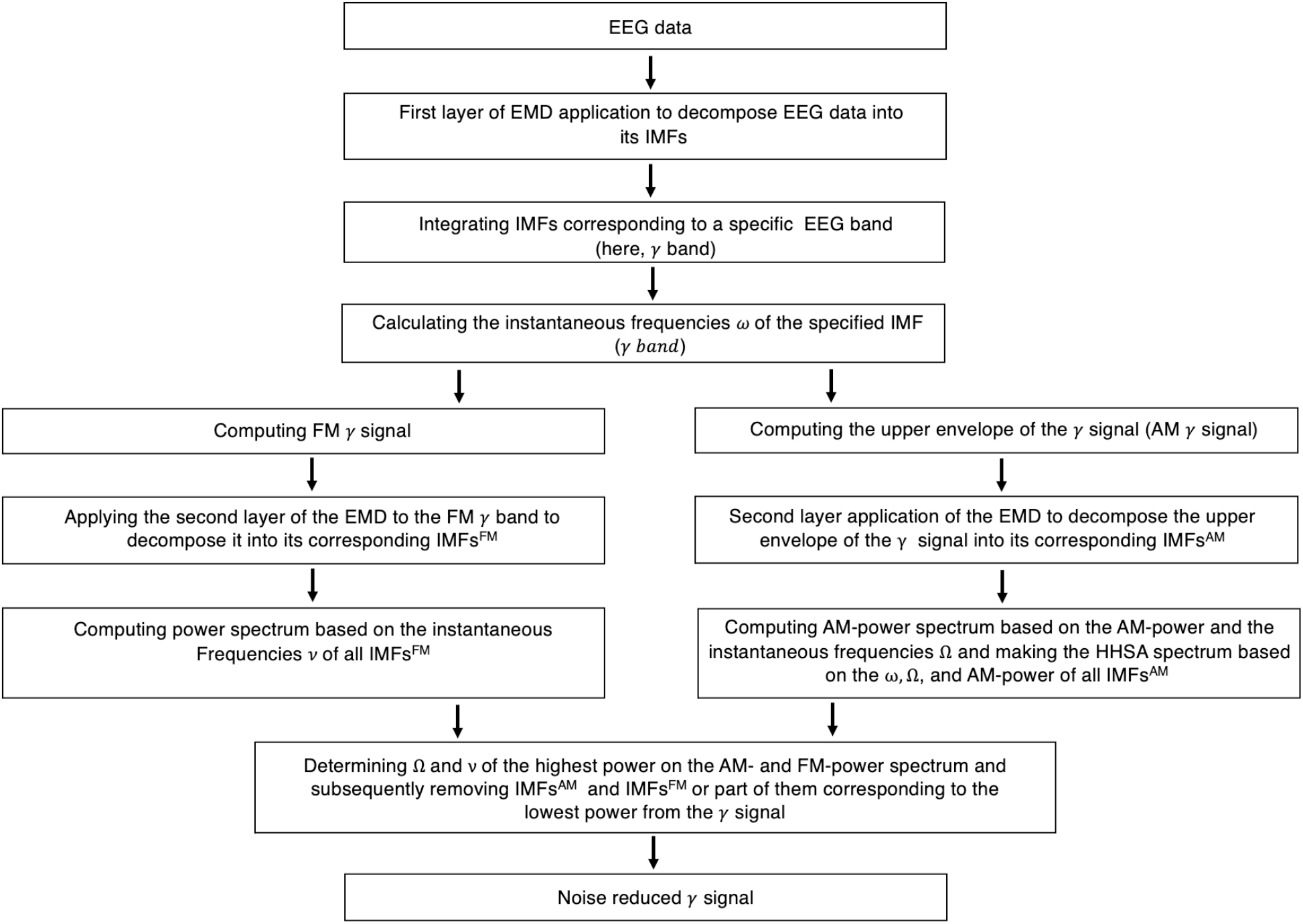
Signal-processing flowchart of the proposed approach.

As mentioned above, when an IMF contains desired frequencies with high energy and undesired frequencies with low energy, only a part of an IMF can be removed. These types of IMFs^FM^ and IMFs^AM^ can be identified by their power spectrum. Part of an IMF is removed by zeroing all frequencies other than the frequency range of interest. This usually happens in the IMFs corresponding to high frequencies in both the AM and FM spectrums. It should also be stressed here that for denoising the FM component of a signal, two cases are possible in the FM power spectrum: 1) a single peak and 2) mode-mixing or multiple peaks. When a signal contains components of close frequencies with almost the same amplitude, there might be more than one frequency in some of its IMFs after decomposition, which is typical [53, 54]. In the single-peak case, IMFs^FM^ with frequencies in the peak range of the FM-power spectrum are kept, and IMFs^FM^ that do not have high power are removed from the FM part of the signal. In the multiple-peak case, just IMFs^FM^ with very low powers are removed, so IMFs^FM^ over a broader range of frequencies are maintained compared to single-peak case. This is because IMFs^FM^ with not very low power might be created by frequency modulation in the original signal. The frequency range of the frequency-modulated signal varies between above and below the carrier frequency, depending on the maximum modulating frequency in the signal. By including more IMFs^FM^ in the FM part of the signal comprised of more than one frequency, losing part of the original signal and its information is prevented. To investigate the similarity between the pure signal (simulated signal before adding noise) and the denoised signal computed by the HHSA method and the proposed approach, we computed Mutual Information (MI) [55] using Neuroscience Information Theory Toolbox (https://github.com/nmtimme/Neuroscience-Information-Theory-Toolbox).

## III. Results

### A. Simulation

The simulated pure and noisy gamma-band signals and their frequency spectra are respectively shown in Fig. 2A– 2F for signal consisting of a single frequency of 30 Hz with SNR of *≈*2 dB (*s-signal1*), and in Fig. 3A–3F for signal containing 30 Hz, 60 Hz and 75 Hz with SNR of *≈* 6 dB (*ssignal4*). Signals with two different SNR values selected just for illustration. After computing the upper envelopes of the noisy signals, Figs. 2G and 3G, they are further decomposed into IMFs^AM^ using an EMD-based method, ICEEMDAN, with 300 ensembles, i.e., number of copies of the input signal, and a level of noise of 0.2 [56].

**Fig. 2:**
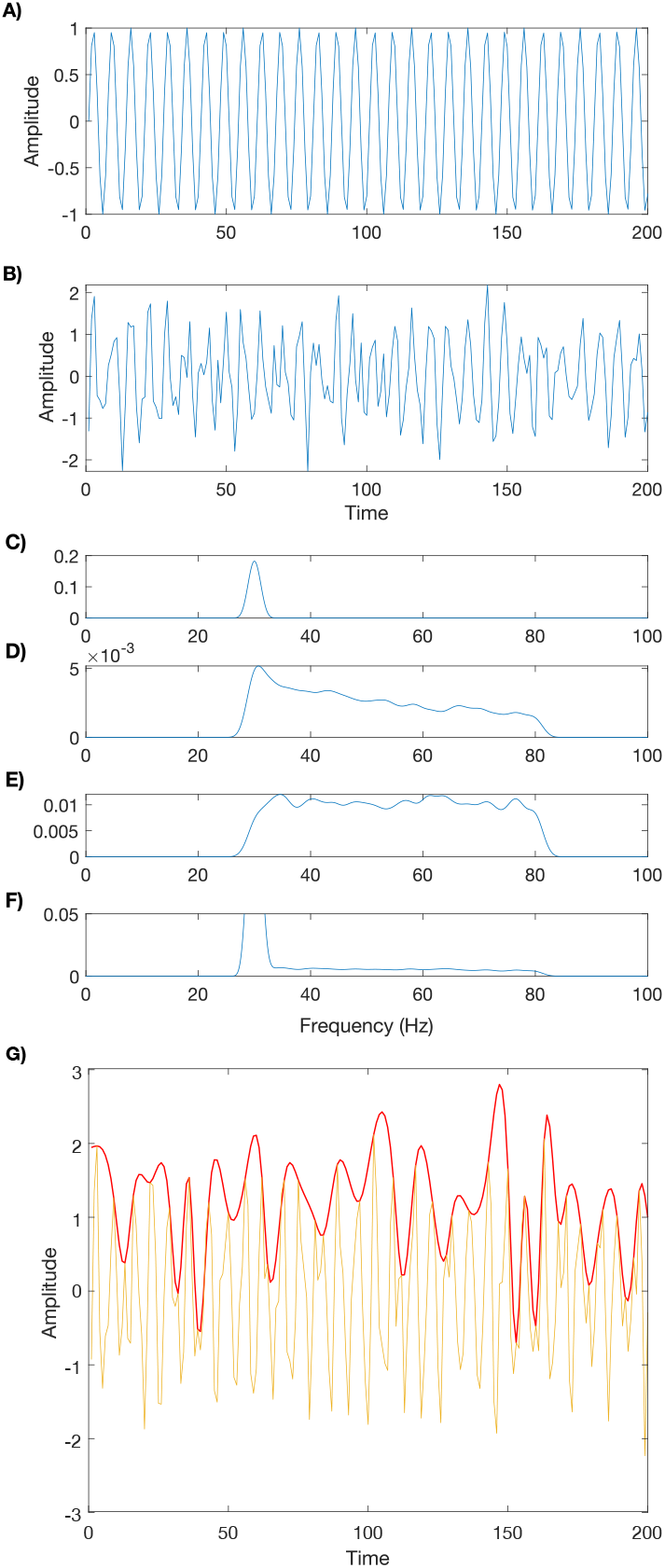
Representation of the simulated pure and noisy signals with carrier frequency 30 Hz, their power spectrum and the upper envelope of the noisy signal. A) Simulated pure and B) the noisy signals. 200 time points are shown for illustration. Power spectrum of C) pure signal, D) filtered pink noise, E) filtered white noise, and F) noisy signal. G) Upper envelope of the simulated noisy signal.

**Fig. 3:**
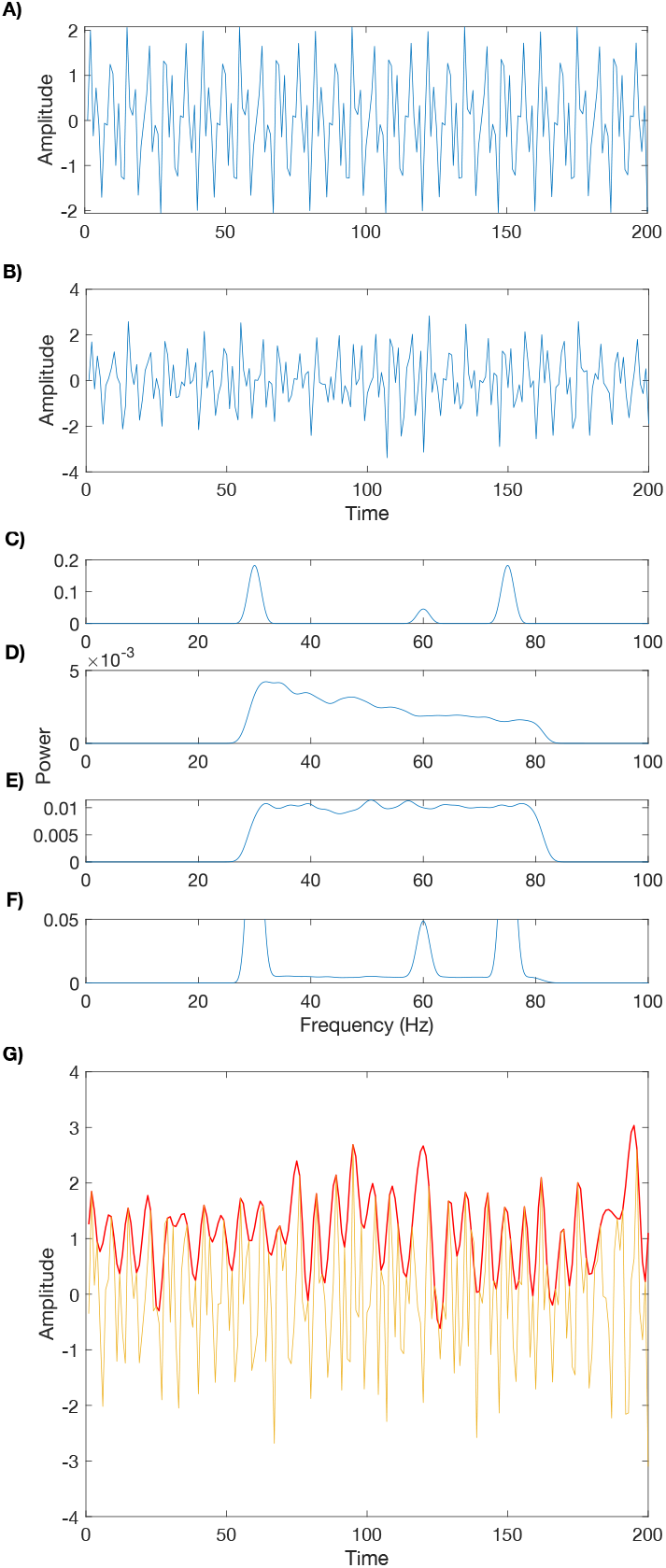
Representation of the simulated pure and noisy signals containing 30, 60, and 70 Hz frequencies, their power spectrum and the upper envelope of the noisy signal. A) Simulated pure and B) the noisy signal. 200 time points are shown for illustration. Power spectrum of C) pure signal, D) filtered pink noise, E) filtered white noise, and F) noisy signal. G) Upper envelope of the simulated noisy signal.

According to the 2D power spectrum and the HHSA spectrum of the upper envelope (steps 5 and 7 in §II-C), the envelope-modulation frequencies are less than *≈*17–20 Hz for *s-signal1* (see Figs. 4A and 4B) and between 20 to 35 Hz for *s-signal4* (see Figs. 5A–5B) where the power density is highest. Thus, as stated in step 8 of the proposed method, IMFs^AM^ (shown in Figs. 4C and 5C)) with frequencies other than these frequency ranges are removed from each signal’s upper envelope. Accordingly, to denoise the AM component of *s-signal1*, IMF^AM^1 and IMF^AM^10 to IMF^AM^11 were removed. Frequencies higher than 20 Hz in IMF^AM^2 were also zeroed. To denoise the AM component of *s-signal4*, IMF^AM^3 to IMF^AM^11 were excluded based on the power spectra shown in Figs. 5A and 5B. Furthermore, since white noise is a wideband signal without modulation, it has a modulation frequency of *≈* 0 Hz. Thus, IMFs^AM^ with a frequency lower than 0.1 Hz were also removed for both simulated signals except for the last IMF^AM^ that represents the input signal’s trend [7].

**Fig. 4:**
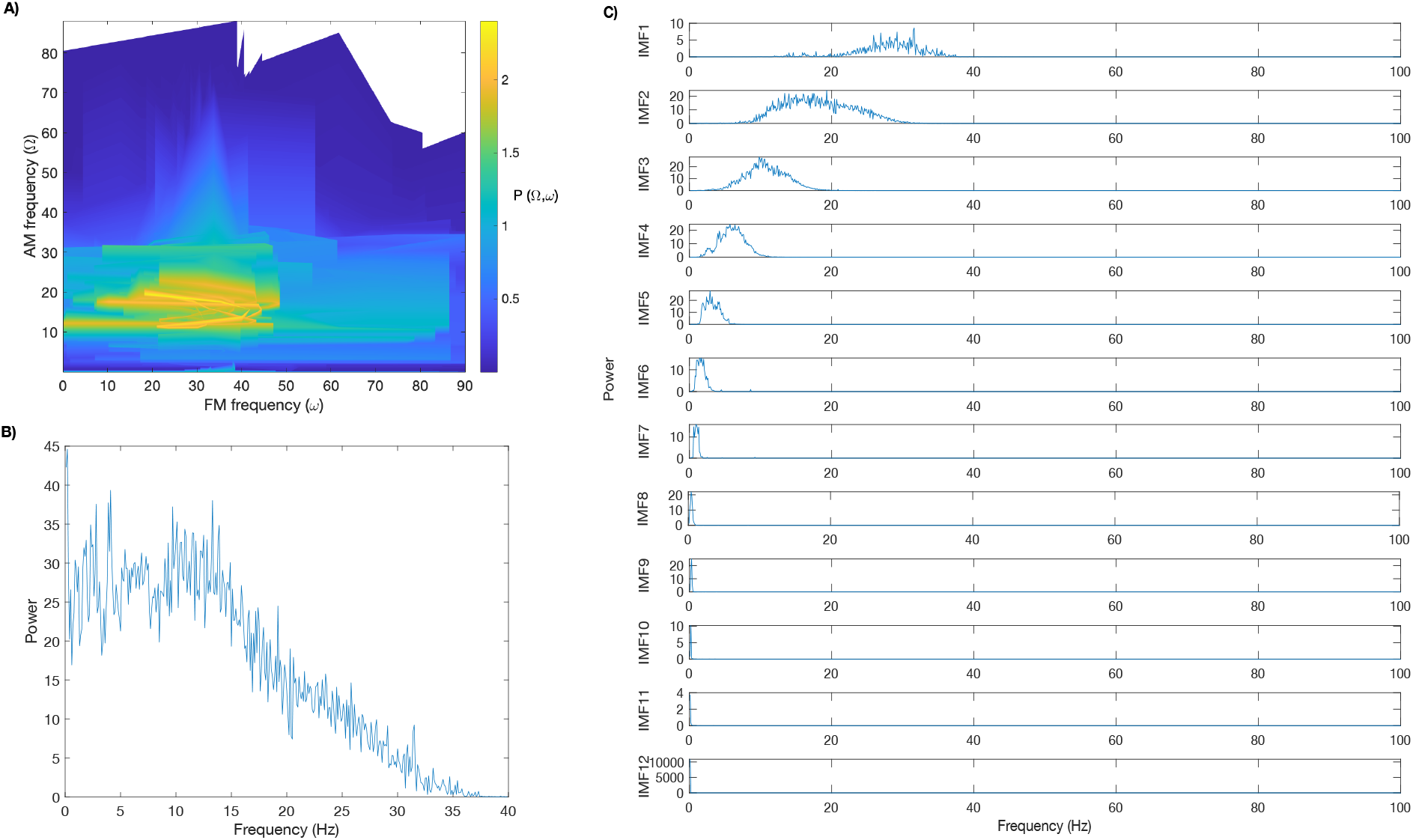
HHSA spectrum of the noisy signal with carrier frequencies of 30 Hz, its AM power spectrum, and the power spectrum of the IMFs^AM^ of the noisy signal. A) HHSA spectrum of the noisy signal computed using the HHT method, which is based on the instantaneous power and frequencies of all IMFs^AM^ and signal’s FM instantaneous frequencies. B) Power spectra based on the instantaneous frequencies of all IMFs^AM^. C) Power spectrum of each of the IMFs^AM^.

**Fig. 5:**
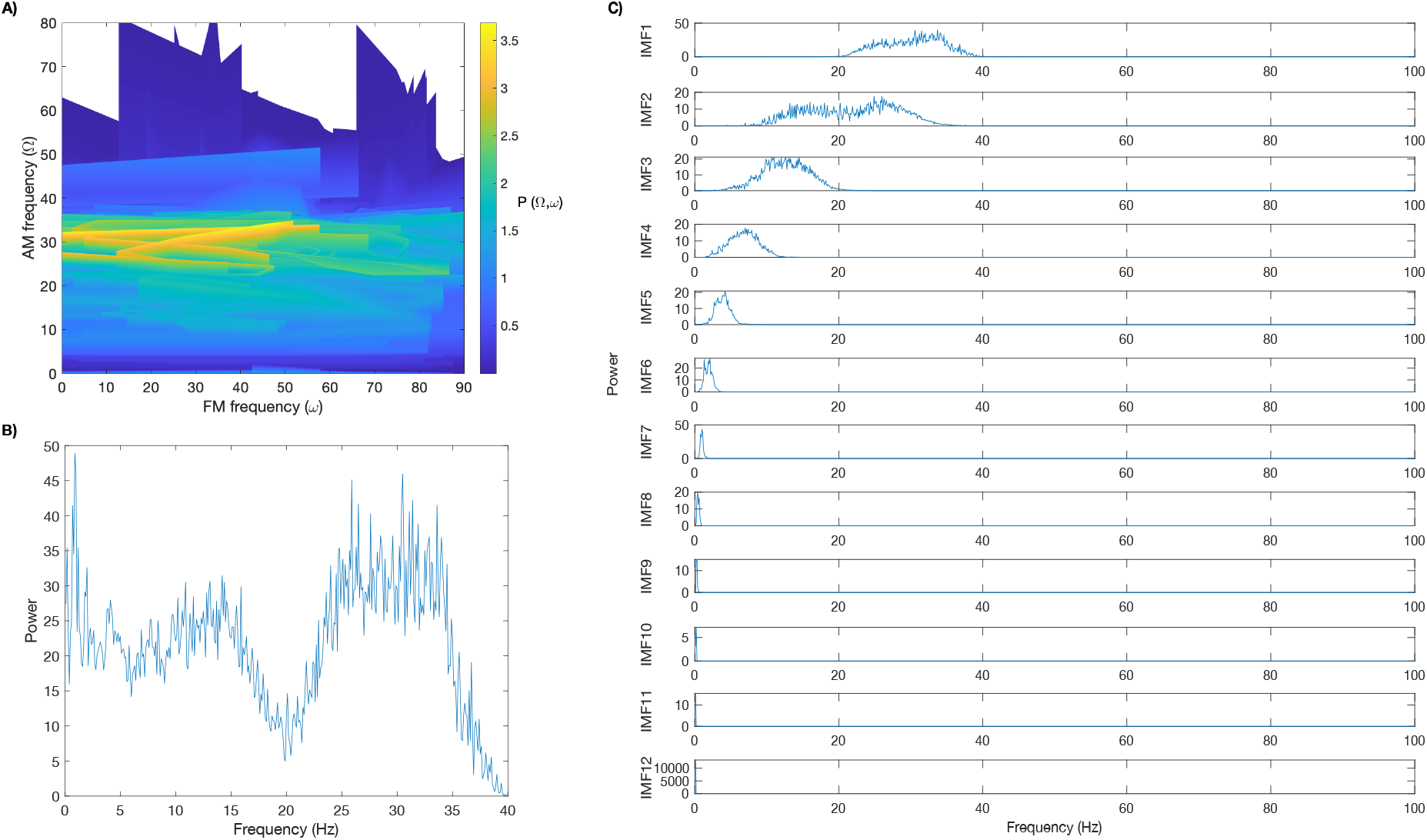
HHSA spectrum of the noisy signal with combination of three frequencies 30, 60, 75 Hz), its AM power spectrum, and the power spectrum of the IMFs^AM^ of the noisy signal. A) HHSA spectrum of the noisy signal based on the instantaneous power and frequencies of all IMFs^AM^ and signal’s FM instantaneous frequencies. B) Power spectra based on the instantaneous frequencies of all IMFs^AM^. C) Power spectrum of the IMFs^AM^.

To denoise the FM part of *s-signal1* and *s-signal4*, according to step 12 in §II-C, the power spectrum of the FM component and its IMFs^FM^ are computed, illustrated in Fig. 6A–6D. According to step 13 of the method, to denoise the FM component of *s-signal1*, IMF^FM^3 to IMF^FM^14 (last IMF), shown in in Fig. 6B, as well as frequencies higher than 35 Hz in IMF^FM^1 were removed. For denoising the FM component of *s-signal4*, IMF^FM^8 to IMF^FM^13 (last IMF) that contains frequencies with the lowest power, depicted in Fig. 6D, were removed from the FM component of the *s-signal4*.

**Fig. 6:**
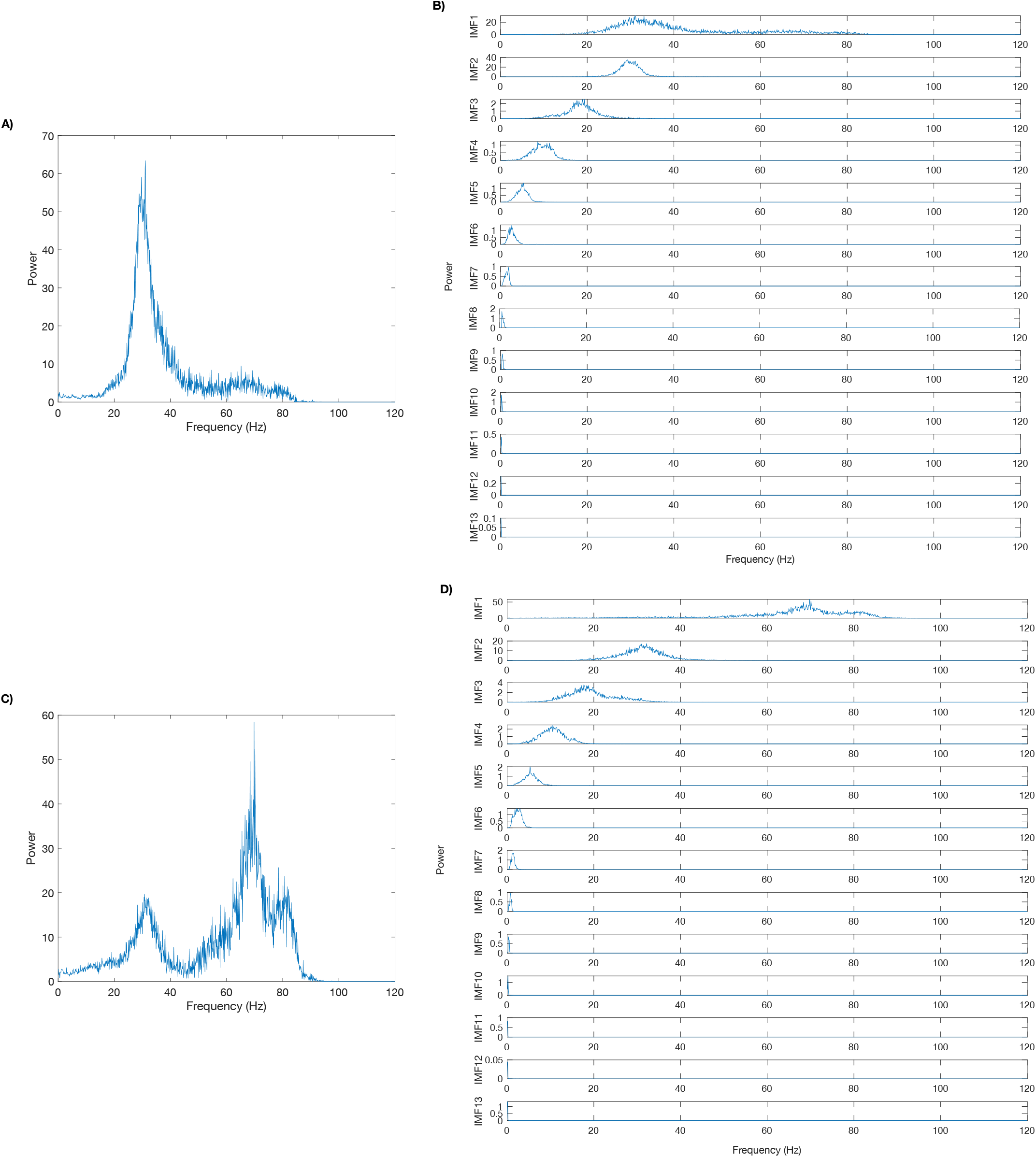
Power spectrum of the frequency-modulated noisy signals and their IMFs^FM^. A) and B) are respectively power spectrum of the FM component of the 30-Hz signal and its IMFs^FM^. C) and D) are respectively power spectrum of the FM component of the signal made of frequencies 30, 60 and 75-Hz, and its IMFs^FM^.

It should be noted that, as seen in the simulations, to denoise the AM component in both simulation cases, just IMFs^AM^ of frequencies within the range of frequencies with the highest power in the power spectrum (i.e., frequencies in the bandwidth) are maintained (Figs. 4C and 5C). However, to denoise the FM component of *s-signal4*, a signal with more than one prominent peak in its FM-power spectrum, shown in Fig. 6C, only the very low-power IMFs^FM^ are removed. As explained in the method section, having more than one peak in the power spectrum represents the possibility that some frequencies that are not of very high power are caused by the combination of signals with different frequencies and contain information. In this situation, deciding which frequencies should be removed is risky, and it is better to include more frequencies with the cost of having more noises rather than losing information.

Figure 7 demonstrates the comparison between the simulated pure signal and the resultant denoised signal using the HHSA method and our method for *s-signal1*, and Fig. 8 demonstrates the same comparison for *s-signal4*.

**Fig. 7:**
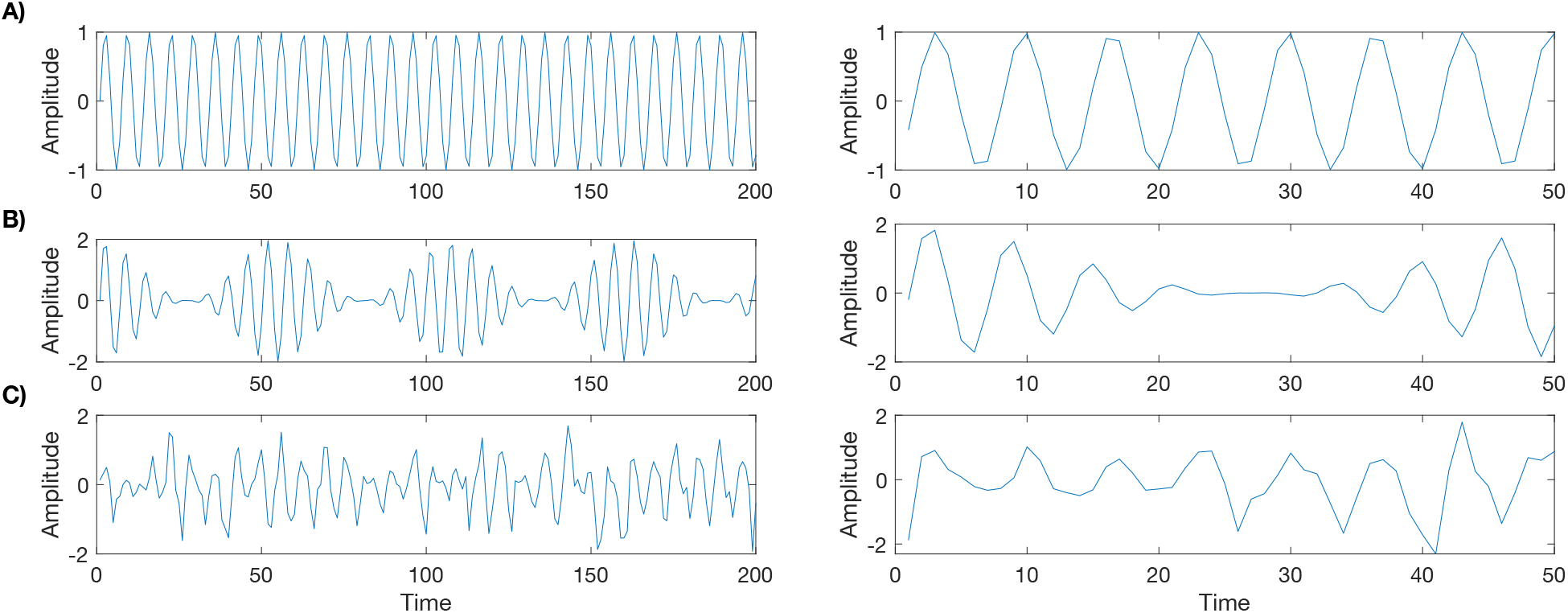
Comparison between the pure signal with frequency 30 Hz, the signal denoised by the HHSA and our methods. Here, 200 time points of the signals are shown for illustration. A) Pure signal. B) Denoised signal by the HHSA method. C) Denoised signal by the proposed method. The right panel shows 50 time points of each of the signals in the left panel just for illustration.

**Fig. 8:**
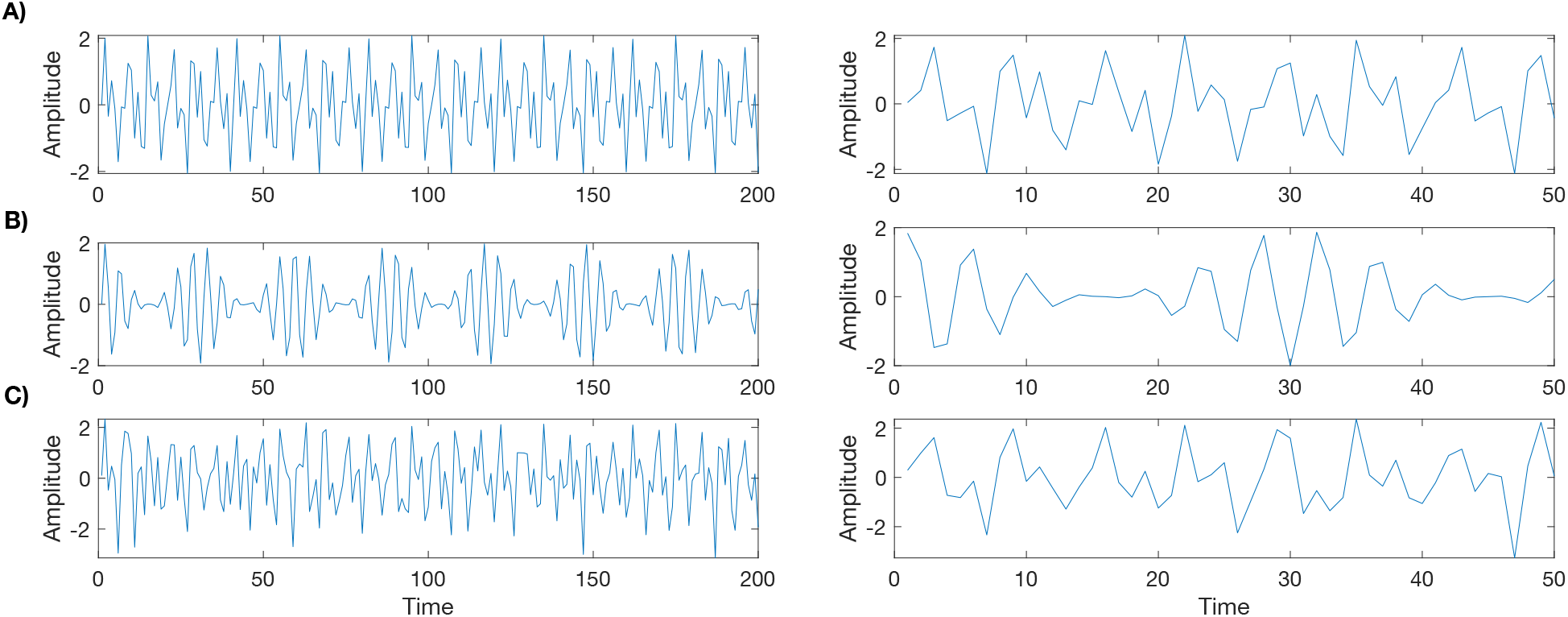
Comparison between the pure signal, the signal denoised by the HHSA and our methods when it is made of a combination of 30, 60, and 75 Hz frequencies. A) Pure signal. B) Denoised signal by the HHSA method. C) Denoised signal by the proposed method. The right panel shows 50 time points of each of the signals in the left panel just for illustration.

As shown in Figs. 7C and 8C, our proposed method identifies the pure signal’s peaks, depicted in Figs. 7A and 8A, better than the HHSA method, shown in Figs. 7B and 8B. Table I shows the normalized MI computation results for the HHSA and the proposed method applied to signals with different levels of SNR and PSNR. We generated 1000 realizations of each signal to find the frequencies with significant power. The table includes the Ω and *ω* computed through the HHSA method with p-value *≤*0.001 and also the IMFs^FM^ and IMFs^AM^ need to be removed in the proposed method. Odd-numbered signals are made of just one carrier frequency, and even-numbered signals are made of a combination of three career frequencies, as explained in §II-B.

**TABLE I:**
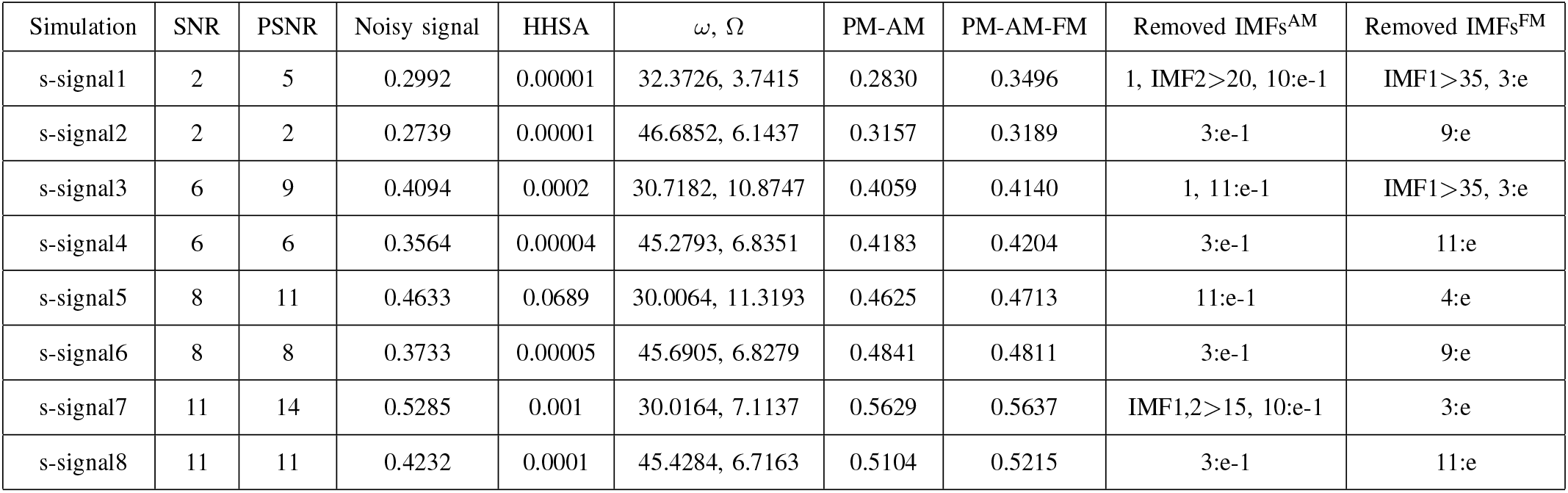
Comparison of the MI between the simulated pure signal and its denoised version using the HHSA and the proposed method (p-value*≤* 0.001). Odd- and even-numbered signals are made of one and a combination of three career frequencies, respectively. Ω and *ω* are computed by the HHSA method; IMFs^FM^ and IMFs^AM^ are the removed IMFs in our method. The column “Noisy signal” shows the MI between the simulated pure (without added noise) and the noisy signals. PM is referred to “proposed method”. Columns PM-AM and PM-AM-FM show the MI between the pure signal and the denoised signal using our proposed method when just the amplitude is denoised and when both the amplitude and phase are denoised, respectively. Here “e” means the “end IMF” or the final IMF, e.g., “3:e” means IMF^AM^3 to final IMF^AM^ are removed from the AM part.

The column “Noisy signal” shows the MI between the simulated pure (without added noise) and the noisy signals. Columns “PM-AM” and “PM-AM-FM” show the MI between the pure signal and the denoised signal using our proposed method when just the amplitude component is denoised and when both the amplitude and phase of the signal are denoised, respectively. The higher the MI value, the more mutual information and similarities are shared between the two signals. Results of our method, presented in columns “PM-AM” and “PM-AM-FM” of table I, show higher MI between the pure and denoised signals compared to the HHSA method and compared to the case where no denoising is applied (column “Noisy signal” of table I).

### B. Real data

To show our method’s performance on real EEG data (not simulated), we applied the method to a sample of real EEG signal recorded during an EEG-fMRI experiment (see §II-A). As shown in Fig. 9, the EEG signal was decomposed into its IMFs applying the ICEEMDAN method [56], with 300 ensembles and a level of noise of 0.2 to have an optimal decomposition of the EEG signal. The frequency spectra of these IMFs corresponded to the EEG frequency bands used in the literature as follows: summation of IMF1 and IMF2 (gamma,*>* 30 Hz), IMF3 (beta, 12–30 Hz), IMF4 (alpha, 8– 12 Hz), IMF5 (theta, 4–8 Hz), and the summation of IMF6 to IMF18 (delta, 0.1–4 Hz). Figs. 10 and 11 represent the overview of our method applied to the real gamma-band.

**Fig. 9:**
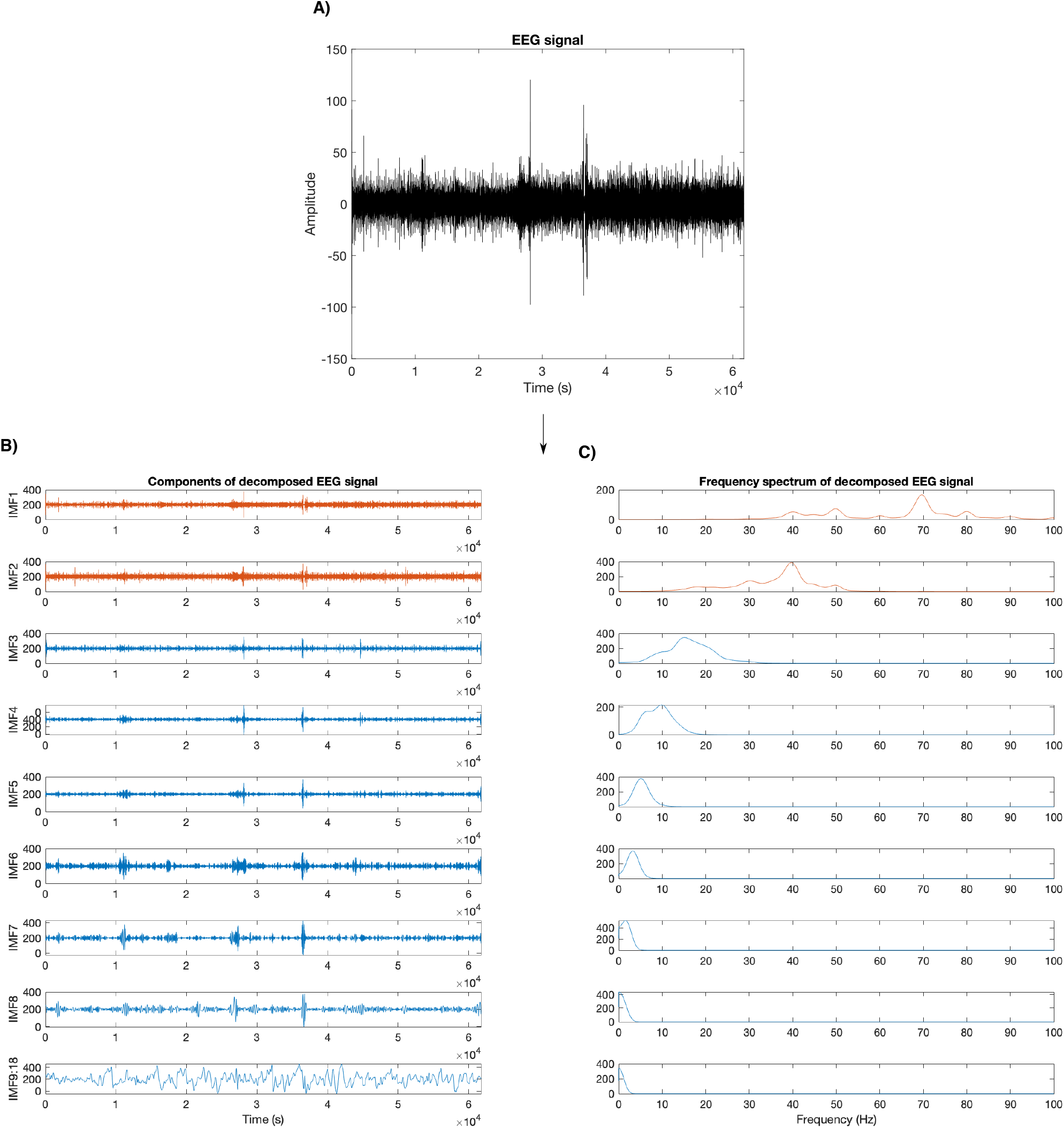
EEG decomposition into its different bands. A) A sample of EEG signal is decomposed by applying ICEEMDAN method to specify the gamma band using frequency spectrum. B) IMFs of the EEG signal. C) IMFs’ frequency spectrum. Here, gamma-band signal is made by the summation of IMF1 and IMF2.

**Fig. 10:**
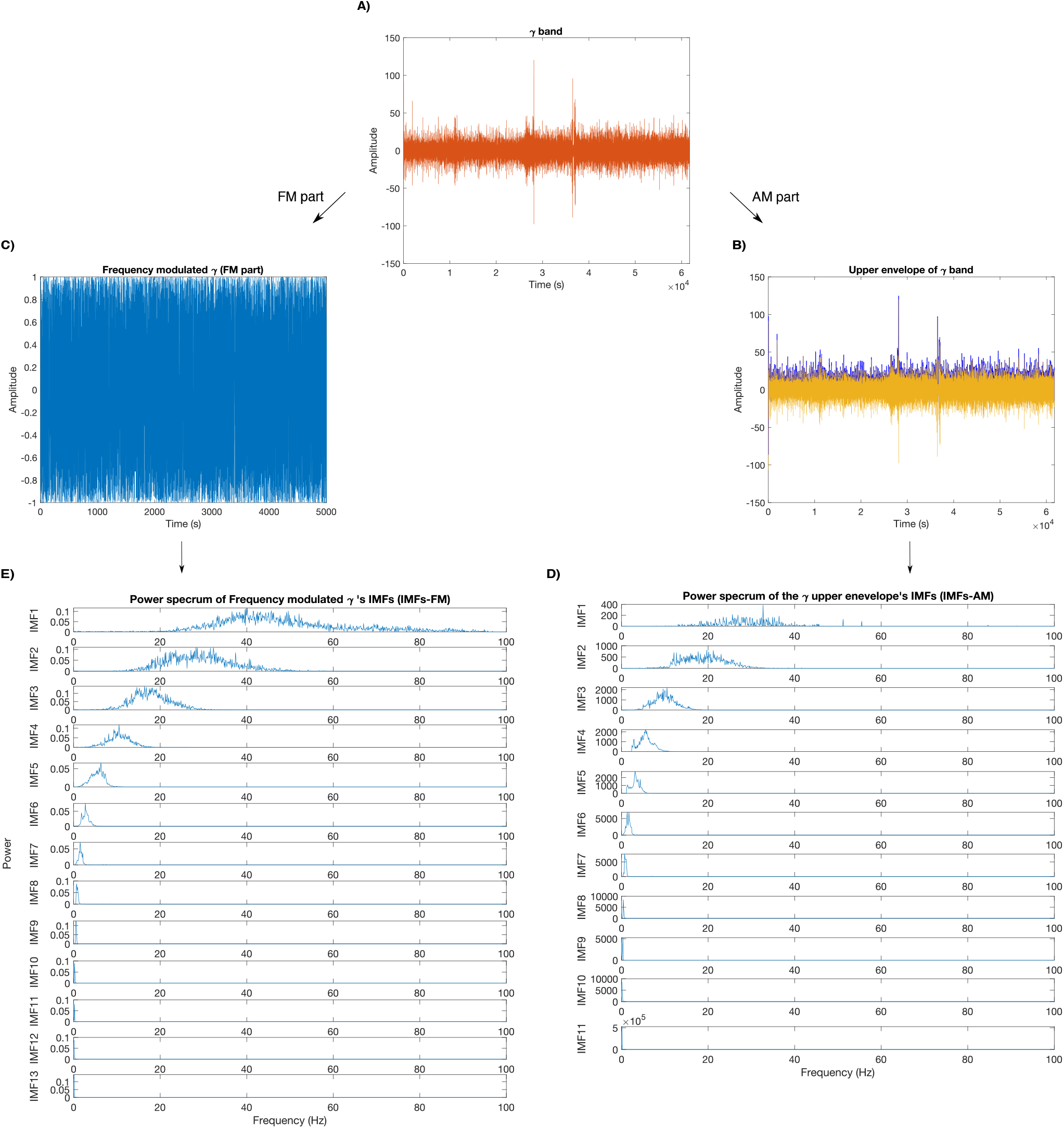
Computing the AM and FM components of the real gamma-band and their power spectrums. A) gamma-band signal, B) the upper envelope of the gamma signal and C) the frequency modulated gamma signal. D) and E) power spectrums of the decomposed AM and FM components, (IMFs^AM^ and IMFs^FM^), respectively.

**Fig. 11:**
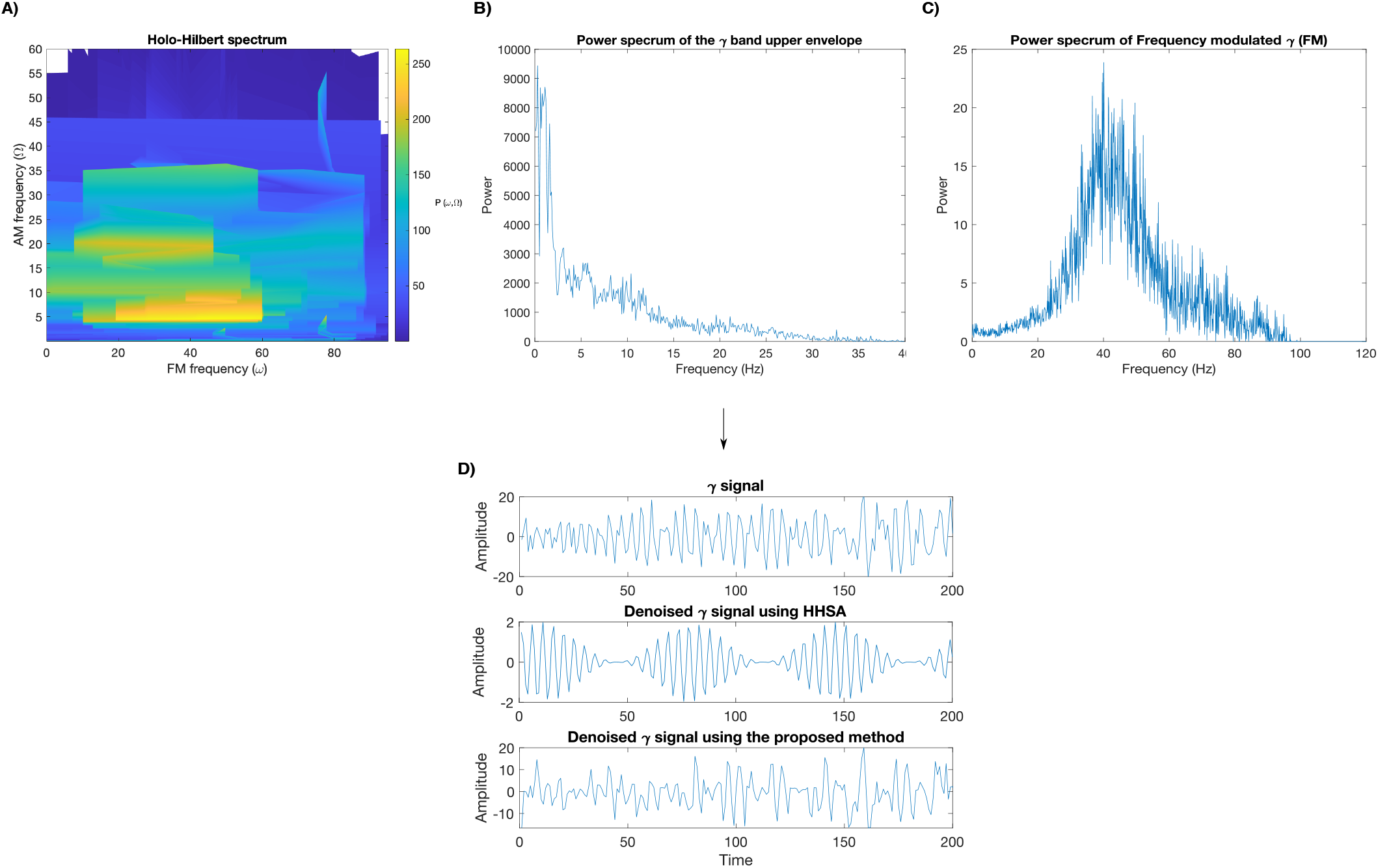
The HHSA plot and the 2D power spectrum of the AM and FM components of the real gamma-signal. A) The HHSA power spectrum. B) and C) The power spectrum of the AM and FM parts, respectively. Denoising the upper envelope by removing IMF^AM^1 and IMF^AM^4 to IMF^AM^10 and also removing frequency higher than 11 Hz from IMF^AM^2 based on the power spectrum of the gamma signal’s upper envelope in (B). FM part of the signal was denoised by removing IMF^FM^3 to IMF^FM^13 based on its power spectrum in (C). Furthermore, frequencies higher than 55 Hz were removed from IMF^FM^1. Comparing the noisy gamma signals with its different denoised versions.

After constructing the gamma band by summing IMF1 and IMF2 of the real EEG signal, Fig. 10A, the upper envelope was computed, Fig. 10B, and decomposed into IMFs^AM^. Based on the power spectrum of the IMFs^AM^, see Fig. 10D, the AM components of the gamma signal were denoised by removing IMF^AM^1 and IMF^AM^4 to IMF^AM^10, in which the power intensity was low. Afterwards, the IMFs^AM^ were further denoised by removing IMF’s frequencies (not a whole IMF) that are higher and lower than the frequencies of the peak’s valley points identified in the 2D power spectrum of gamma signal’s upper envelope, see Fig. 11B. In this case, frequencies higher than 11 Hz were removed from IMF^AM^2.

On the other hand, the signal’s FM portion,shown in Fig. 10C, was denoised by removing IMF^FM^3 to IMF^FM^13, as IMF^FM^1 and IMF^FM^2 had the highest power in frequencies between *≈*25 Hz to *≈*55 Hz, which are the peak’s valley points in the FM power spectrum, see Figs. 10E and 11C. Furthermore, frequencies higher than 55 Hz were removed from IMF^FM^1 because of the valley points of the peak at *≈*42 Hz in the FM power spectrum, Fig. 11C. Applying the HHSA method to the real EEG data obtained the carrier frequency of *ω≈* 41.4607 and AM frequency of Ω *≈*2.2527 Hz, which are in accordance with the area of high power density in the HHSA plot indicated in Fig. 11A. Fig. 11D, shows the comparison of the noisy gamma signal with its different denoised versions computed by the HHSA and our proposed methods.

Figure 12 compares the power spectral density of the denoised real gamma signal computed by the HHSA and the proposed method. Denoised gamma signal computed by the proposed method shows almost the same power density as the noisy signal. The proposed method causes a slightly lower power density in high frequencies, which might be because of the EMG- and MR-related noises that are mainly in high-frequency EEG signals. Thus, lower power density after applying our method might be in consequence of removing these artifacts from the signal that had not been adequately denoised by the pre-existing methods during preprocessing of the data. However, the HHSA method obtained a completely different power spectrum density for the denoised signal, except for the high power density for frequencies *≈* 40 Hz.

**Fig. 12:**
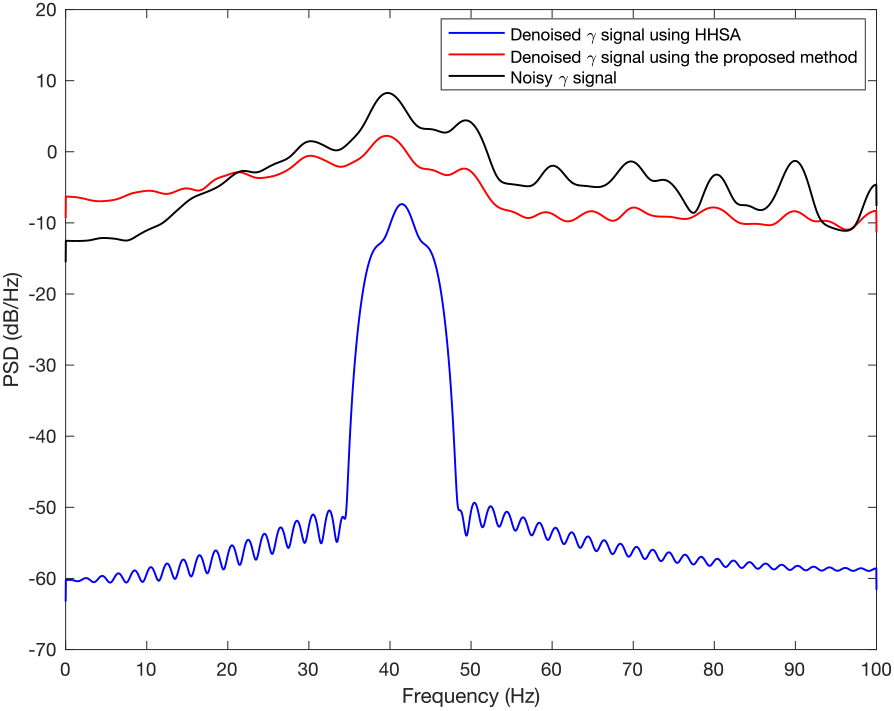
Comparison of the power spectrum density for the HHSA and our method applied to a sample real gamma signal.

## IV. Discussion

In this study, we proposed a method for denoising the EEG gamma band. Our proposed method is based on the Holo-Hilbert method and denoises any nonstationary and nonlinear time-series data. However, in this paper, we specifically focused on gamma-band noise suppression due to its crucial role in mapping the brain’s function.

Our result showed that the HHSA method does not perform well for cases with more than one peak in the FM-power spectrum of a signal compared to the case where a single frequency exists in each IMFs after the first application of the EMD. Each IMF computed by the EMD method is typically assumed to contain only one frequency, but this is seldom the case [53, 54] as in our second simulated gamma-band signal. For cases with multiple peaks in the FM-power spectrum, the HHSA method goes through multiple iterations and applies the EMD method repetitively until no more cyclic characteristics are in the decomposed signal. Consequently, the HHSA method is not computationally efficient. Furthermore, there could be no clear discrepancy between the power peaks in the computed HHSA spectrum. In this situation, different frequency bands cannot be specified in the HHSA spectrum. Consequently, the HHSA method results in wrong frequencies for the AM and FM components.

However, our proposed method does not have these problems as it applies just two layers of EMD to the AM and FM components of the signal. Besides, the denoised signal in our method is obtained differently and is based on the remaining IMFs after removing noisy IMFs^AM^ and IMFs^FM^. Our method yields higher MI between the pure and denoised signals compared to the HHSA method and when the signal is not denoised. Higher MI confirms that our approach extracts the pure signal from the noisy signal with higher efficiency. Moreover, our method yields lower power density in high frequencies of the denoised real gamma signal compared to its noisy version. Lower power in high frequencies could be evidence for our method’s potential to detect and remove problematic artifacts in high-frequency EEG signals, notably gradient artifacts due to abrupt head motion, which are mostly in high frequencies [10, 31, 32].

It is worthy to note that, unlike conventional predefined basis methods, e.g., Fourier analysis and wavelet methods [9], the AM and FM components are separately decomposed in our proposed method. Accordingly, the AM and FM frequencies are detected separately, which reduces the uncertainty of FM-frequency detection and therefore increases the accuracy of a spectral analysis method [7]. Additionally, our proposed method does not require any reference signal or any assumption on the noisy EEG data and is applicable to all types of data (stationary and nonstationary, linear and nonlinear).

We now discuss the limitations of our method. First, the IMFs^FM^ and IMFs^AM^ required to be removed from the signal vary from subject to subject. Second, these IMFs are specified approximately based on their power spectrum during the noise-reduction process. The difference between powers of successive IMFs could be quite small. In this case, constructing a decision boundary for removing or maintaining an IMF becomes challenging. This argument also holds for choosing the frequencies that need to be removed from a specific IMF based on the 2D power spectrum. Furthermore, we used simulated data instead of real EEG signals to check the proposed method’s efficacy as denoised EEG signal is unknown in a real EEG signal. Our method, therefore, has the general limitations of using simulated EEG signal versus real signal.

To conclude, our proposed method efficiently reduces noise from the EEG gamma band by analyzing its AM and FM components. This method applies two layers of EMD to adaptively decompose a signal into IMFs with the signal’s time-varying amplitude and frequency features. Then the IMFs^AM^ and IMFs^FM^ (or parts of them) with low powers are removed from the signal’s AM and FM components as they correspond to noise. Our method would improve the interpretation and localization of the underlying brain’s active regions by providing an efficiently denoised gamma signal. The proposed method could also lead to a better realization of neurovascular coupling and brain function in studies that use simultaneous EEG/fMRI data with more problematic gammaband denoising.

## Acknowledgement

This work was supported by grant number RGPIN-2015- 05966 from Natural Sciences and Engineering Research Council of Canada.

All computations are performed in Matlab. For convenience and replicability, the Matlab code used for this study is provided at https://github.com/nargesmoradi/EEG_denoising_method/tree/main.

## Authors’ Contribution

NM and RS designed the research. NM simulated and analyzed the data, interpreted the results and wrote the manuscript. PL acquired and preprocessed real EEG data and helped with manuscript evaluation. BA acquired and preprocessed real EEG data. BG helped with data interpretation and manuscript evaluation. RS supervised the development of the work, helped with data interpretation and manuscript evaluation.

